# Transcription factor prediction using protein 3D secondary structures

**DOI:** 10.1101/2024.03.14.585054

**Authors:** Jeanine Liebold, Fabian Neuhaus, Janina Geiser, Stefan Kurtz, Jan Baumbach, Khalique Newaz

## Abstract

**Motivation:** Transcription factors (TFs) are DNA-binding proteins that regulate gene expression. Traditional methods predict a protein as a TF if the protein contains any DNA-binding domains (DBDs) of known TFs. However, this approach fails to identify a novel TF that does not contain any known DBDs. Recently proposed TF prediction methods do not rely on DBDs. Such methods use features of protein sequences to train a machine learning model, and then use the trained model to predict whether a protein is a TF or not. Because the 3-dimensional (3D) structure of a protein captures more information than its sequence, using 3D protein structures will likely allow for more accurate prediction of novel TFs.

**Results:** We propose a deep learning-based TF prediction method *(StrucTFactor)*, which is the first method to utilize 3D secondary structural information of proteins. We compare StrucTFactor with recent state-of-the-art TF prediction methods based on *∼*525 000 proteins across 12 datasets, capturing different aspects of data bias (including sequence redundancy) possibly influencing a method’s performance. We find that StrucTFactor significantly (*p*-value *<* 0.001) outperforms the existing TF prediction methods, improving the performance over its closest competitor by up to 17% based on Matthews correlation coefficient.

**Availability:** Data and source code are available at https://github.com/lieboldj/StrucTFactor and on our website at https://apps.cosy.bio/StrucTFactor/

**Supplementary information:** Included

## 1 Introduction

### 1.1. Background and motivation

Transcription factors (TFs) are DNA-binding proteins that influence the transcription regulations of genes (Latchman, 1997). Thus, knowing which proteins are TFs would help to better understand gene regulatory systems (Lambert et al., 2018). Because the identification of TFs using wet-lab experiments is costly and time-consuming (Geertz and Maerkl, 2010), computational methods for predicting TFs have been developed (Kummerfeld and Teichmann, 2006; Wilson et al., 2007; Eichner et al., 2013; Zheng et al., 2008; Kim et al., 2021; Du et al., 2023; Ledesma-Dominguez et al., 2024). Traditional computational methods predict a protein as a TF if the protein shows high sequence similarity to any of the known TF-related DNA-binding domains (DBDs) (Kummerfeld and Teichmann, 2006; Wilson et al., 2007). Because such traditional methods would fail to identify a TF that does not show sequence similarity to any of the known DBDs, data-driven TF prediction methods have been proposed that do not rely on known DBDs (Eichner et al., 2013; Zheng et al., 2008; Kim et al., 2021; Du et al., 2023; Ledesma-Dominguez et al., 2024). Such data-driven methods use features of proteins with known TF labels (i.e., whether a protein is a TF or not) to train a machine learning (ML) model in a supervised manner. Then, the features of a protein (with an unknown TF label) are used as input to the trained model to predict whether the protein is a TF or not.

Existing data-driven methods can be grouped into two categories based on how they extract features of proteins. One category of methods relies on manually-derived protein features. An existing method of this category is TFpredict (Eichner et al., 2013). TFpredict aligns the sequence of a protein to known TFs and non-TFs using the Basic Local Alignment Search Tool (BLAST) (Altschul et al., 1990), and uses the distribution of the corresponding alignment scores as features of the protein. Next, such features of proteins along with their TF labels are used to train a support vector machine, which is then used to predict whether a given protein with an unknown TF label is a TF or not. TFpredict outperformed previously proposed TF prediction methods (Eichner et al., 2013).

Instead of relying on manually-derived protein features, another category of methods, which is the focus of our study, learns features of proteins using deep learning (DL). Two recent methods in this category are DeepTFactor (Kim et al., 2021) and DeepReg (Ledesma-Dominguez et al., 2024). Both methods first represent a protein sequence in a “one-hot” encoding matrix which, for each sequence position of the protein, indicates one of the 20 amino acids. Given such one-hot encoding matrices of proteins with known TF labels, DeepTFactor and DeepReg employ different DL models. While DeepTFactor relies on convolutional neural networks (CNNs), DeepReg further extends DeepTFactor’s CNN-based model by incorporating a recurrent neural network and an attention mechanism module. DeepTFactor was shown to outperform two previously existing TF prediction methods, i.e., TFpredict and P2TF (Ortet et al., 2012). DeepReg was recently shown to outperform DeepTFactor with respect to some, but not all, performance measures (Ledesma-Dominguez et al., 2024).

All of the existing data-driven TF prediction methods rely on sequence features of proteins. However, proteins (including TFs) perform functions in their 3-dimensional (3D) structural forms. For several other tasks, including protein 3D structural classification (Newaz et al., 2022; Guo et al., 2019; Newaz et al., 2020), it was shown that methods using 3D structural (either secondary or tertiary) features outperform methods that only rely on sequence features. Hence, given the large number of high quality (Akdel et al., 2022) 3D protein structures predicted by recent methods such as AlphaFold2 (Jumper et al., 2021) and ESMFold (Lin et al., 2023), it seems feasible to improve upon the current state-of-the-art TF prediction methods by exploiting 3D structural features of proteins.

### 1.2. Our contributions

We propose a novel DL-based TF prediction method called *StrucTFactor* (Figure 1), which is the first TF prediction method to use 3D secondary structural information of proteins. Given a protein 3D structure, StrucTFactor extracts secondary structural information using the dictionary of protein secondary structure (DSSP) (Kabsch and Sander, 1983). StrucTFactor represents the secondary structural information of a protein in a one-hot encoding matrix similar to DeepTFactor. While DeepTFactor’s one-hot encoding of the protein indicates the amino acid, StrucTFactor’s one-hot encoding indicates the amino acid and the secondary structure (i.e., *α*-helix, *β*-sheet, or coil-turn), at each protein sequence position. Given the TF label for each one-hot encoded protein, StrucTFactor trains a CNN-based DL model, which is then used to predict whether a given protein with an unknown TF label is a TF or not.

**Figure 1:**
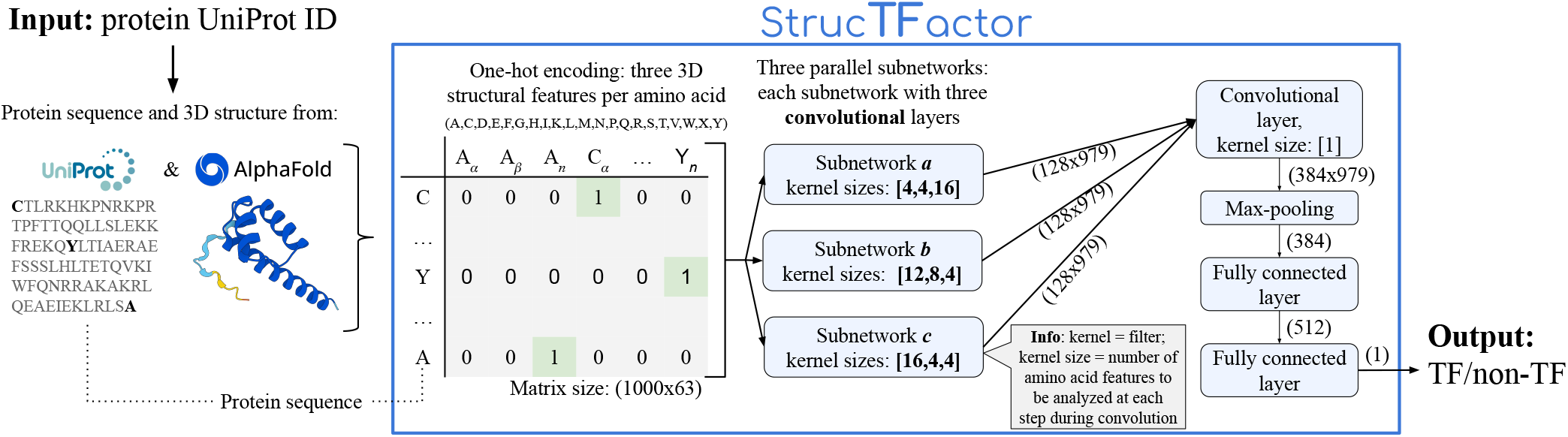
Overview of our TF prediction method: StrucTFactor. See Section 2 for details.

We only compare StrucTFactor with DeepTFactor and DeepReg because these two methods (1) are the most recent TF prediction methods, (2) come with publicly available software implementations, and (3) outperformed previous TF prediction methods. This implies that, if StrucTFactor outperforms DeepTFactor and DeepReg, then it would likely also outperform other previously proposed TF prediction methods. The recently published method TFnet (Du et al., 2023) was not part of our comparison, as there is no publicly available implementation.

In our computational experiments, we use all *∼*525 000 reviewed proteins from UniProt (The UniProt Consortium, 2022) that have their 3D structures in the AlphaFold database (Varadi et al., 2021). Note that, due to limited 3D structural data of TFs in the Protein Data Bank (PDB) (Berman et al., 2000), we could not use PDB data (see Section 2.1 for details). We identify a protein as a TF or non-TF based on an established criterion already used by existing TF prediction methods (Kim et al., 2021; Ledesma-Dominguez et al., 2024), which relies on Gene Ontology (GO) terms (The Gene Ontology Consortium, 2015). Given the *∼*525 000 proteins and their TF labels, we create 12 datasets capturing different aspects of data bias, including sequence redundancy, the numbers of non-TFs vs. TFs (i.e., class assignment ratio), and the quality of 3D protein structure predictions, that can influence the training of an ML model. Given a dataset, we divide it into five non-overlapping subsets while keeping the ratio of the numbers of non-TFs vs. TFs in each subset the same as in the entire dataset. Given a subset, we use it as an *independent* (i.e., not used in training and validation) test data and the remaining four subsets as training and validation data. We do this for each of the five subsets and thus obtain five combinations of training, validation, and independent test data. We evaluate a trained model based on three performance measures, i.e., area under precision-recall curve (AU-PRC), area under receiver operating characteristic curve (AU-ROC), and Matthews correlation coefficient (MCC).

For all 12 datasets, StrucTFactor shows superior performance compared to DeepTFactor and DeepReg. Interestingly, for the most reliable dataset (with sequence non-redundant proteins and the smallest ratio of the numbers of non-TFs vs. TFs), StrucTFactor significantly (*p*-value *<* 0.001) outperforms both DeepTFactor and DeepReg, improving performance over its closest competitor (DeepTFactor) by up to 17% when considering MCC. Using the concept of integrated gradients (Sundararajan et al., 2017), we investigate the parts of a protein that StrucTFactor focuses on to predict the protein as a TF. We find that, in contrast to existing methods, StrucTFactor puts more focus on the known DBDs of TFs. This is interesting because we do not use any DBD information during training of any model. This potentially explains StrucTFactor’s superiority over existing methods, highlighting the importance of considering 3D secondary structural features of proteins to identify novel TFs.

## 2. Data and method

### 2.1. Collection of protein data

We use all 570 830 reviewed proteins from UniProt (last download on 2024-02-22 from https://ftp.uniprot.org/pub/databases/uniprot/current_release/knowledgebase/complete/uniprot_sprot.dat.gz). We only consider a protein further if it (1) has up to 1 000 amino acids, (2) only contains the 20 standard amino acids, and (3) is present in the AlphaFold protein structure database (Varadi et al., 2021) (downloaded 2022-10-19 from https://ftp.ebi.ac.uk/pub/databases/alphafold/latest/swissprot_pdb_v4.tar), resulting in 524 674 proteins. We identify this set of proteins as *allAF*, indicating all eligible proteins from AlphaFold. Note that we do not use 3D protein structures from PDB because of the lack of sufficient 3D structural information about TFs. For example, there are only five TFs with secondary structural information for at least 80% of their residues. For details, see Supplementary Section S1, Supplementary Figure S1, and Supplementary Table S1.

### 2.2 Selection of reliable protein 3D structures

We identify “reliable” 3D protein structures from AlphaFold using an established criterion (Akdel et al., 2022), based on the confidence score provided by AlphaFold for each amino acid of a predicted protein 3D structure. That is, we identify a protein 3D structure to be reliable if and only if *≥* 90% of its amino acids have a confidence score of *≥* 70. Among 524 674 proteins in allAF (Section 2.1), 340 765 are reliable (named *reliableAF*).

### 2.3. Transcription factor annotation

To annotate whether a protein is a TF or not, we rely on an established criterion already used by existing TF prediction methods, i.e., DeepTFactor and DeepReg, that is based on GO terms. We annotate a protein as a TF if it is associated with either (1) a TF GO term or (2) a transcription regulation GO term in combination with a DNA binding GO term, else we annotate the protein as a non-TF. We extract protein-GO term associations from UniProt. We consider the same 21 TF GO Terms, 11 transcription regulation GO terms and 4 DNA-binding GO terms as used by DeepTFactor and DeepReg (Supplementary Table S2).

### 2.4. Identification of sequence non-redundant proteins

We group a set of proteins into sequence non-redundant clusters using the many-against-many sequence searching (MMSeqs2) tool (Steinegger and Söding, 2017). Given a large number of proteins as in our study, MMSeqs2 has been shown to be similarly as accurate as traditional sequence alignment methods such as BLAST (Altschul et al., 1990), while being much more computationally efficient. We run MMSeqs2 using the command: *mmseqs easy-cluster –min-seq-id 0*.*3 -c 0*.*8 –cov-mode 0*. This means that the pairwise sequence identities of proteins within a cluster are *≥*30%, while pairwise sequence identities of proteins across clusters are *<*30%, enforcing that at least 80% of residues from both proteins are included in the alignment. Note that a sequence identity of *<* 30% is a standard proven choice to sufficiently remove sequence redundancy among proteins, which reduces the chances of selecting sequence (and potentially 3D structural) homologous proteins (Rost, 1999). To get a set of sequence non-redundant proteins, we choose one protein per-cluster, as follows. Given a cluster with no TF, we keep the cluster representative protein selected by MMSeqs2. Given a cluster with at least one TF, if the cluster representative protein selected by MMSeqs2 is a TF then we keep it, else we randomly choose one of the TFs from the cluster. We preferably choose a TF from a cluster to keep as much TF data as possible to effectively train and test an ML model.

### 2.5. Creation of datasets

According to the considered TF annotation scheme (Section 2.3) 7 418 of all 340 765 proteins from reliableAF are TFs. Hence, we have a massive (number of non-TFs is *∼*45 times more than the number of TFs) class imbalance problem that can negatively impact the training of an ML model (Megahed et al., 2021). Thus, similarly as done by existing TF prediction methods, i.e., DeepTFactor and DeepReg, we take three random samples from all 333 347 non-TFs, such that the number of non-TFs is either 3, 5, or 10 times more than the number of TFs, resulting in three datasets represented as *D*(*rl, r, s*). Here, *rl* denotes that the protein comes from reliableAF, *r* denotes that the set is potentially sequence redundant, and *s* ∈{3, 5, 10} (Figure 2).

**Figure 2:**
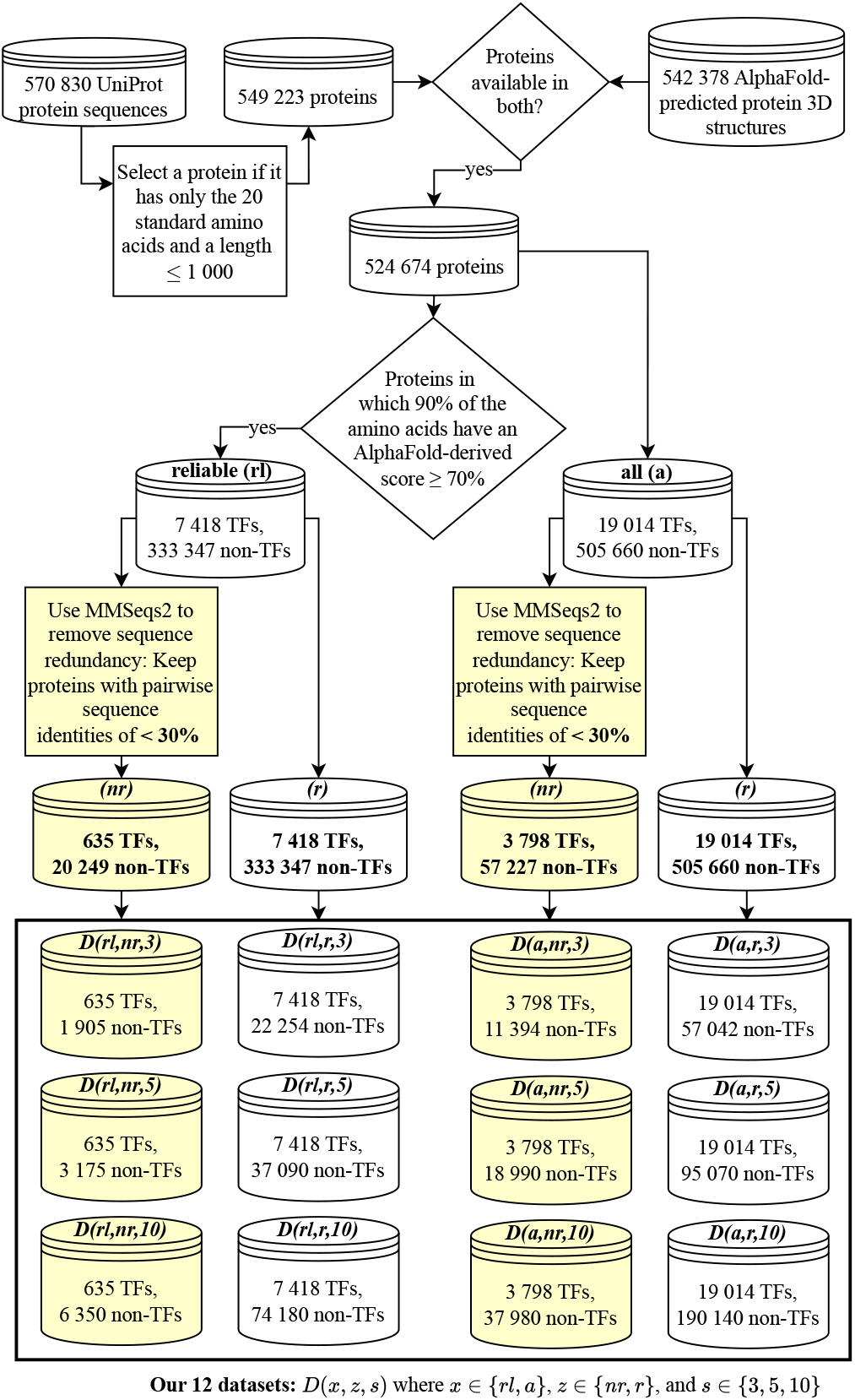
Overview of the creation of our 12 datasets (Section 2.5). We create two sets of proteins, i.e., *allAF* (denoted by the letter code *a*) and *reliableAF* (denoted by *rl*). Among each of allAF and reliableAF protein sets, we create a sequence redundant (letter code *r*) and a sequence non-redundant (letter code *nr*) set of proteins. From each of the sequence redundant and sequence non-redundant protein sets, we create three datasets reflecting the ratio of the numbers of non-TFs vs. TFs of 3, 5, and 10. We name a dataset using the convention *D*(*x, z, s*), where *x* ∈ {*rl, a*}, *z* ∈ {*nr, r*}, and *s* ∈ {3, 5, 10}. In yellow, we highlight the relevant parts of the data creation logic leading to sequence non-redundant datasets.

Sequence redundancy between proteins within and across training and test data could negatively impact the generalization ability of the trained model (Hou et al., 2022; Bernett et al., 2024). Hence, we remove sequence redundancy among the 340 765 reliableAF proteins (Section 2.4), resulting in 20 884 proteins of which 635 are TFs. Given the sequence non-redundant proteins, we handle the class imbalance problem similarly as above resulting in three datasets named *D*(*rl, nr, s*), where *nr* means that the datasets are sequence non-redundant.

Thus, in total, we create six datasets using proteins from reliableAF. Note that because the dataset *D*(*rl, nr*, 3) has sequence non-redundant proteins and the number of non-TFs is only three times greater than the number of TFs, we consider *D*(*rl, nr*, 3) as the most reliable dataset in our study. Analogous to reliableAF, using allAF proteins, we create six datasets denoted as *D*(*a, z, s*), where *a* means that the datasets come from allAF proteins, *z* ∈ {*nr, r*} and *s* ∈ {3, 5, 10} (see Figure 2 and Supplementary Section S2 for details).

### 2.6. Protein features

#### 2.6.1. Existing sequence-based features

DeepTFactor and DeepReg represent a protein of length, say *l*, as a one-hot encoding *𝓁×* |*R*|-matrix *M*_seq_, where *R* is the alphabetically ordered set of the standard 20 amino acids plus a wildcard *X* for all other amino acids. An entry *M*_seq_[*i, j*] is 1 if and only if the amino acid at the *i*^*th*^ position in the protein sequence is the *j*^*th*^ amino acid in *R*.

#### 2.6.2. Our proposed 3D structure-based features

Given the 3D structure of a protein, we extract its secondary structural features using DSSP, which annotates each amino acid of a protein structure as one of the seven secondary structural categories or a placeholder (blank output) to represent an undetermined secondary structural output. We combine the seven secondary structural categories and the blank output obtained from DSSP into three disjoint secondary structural groups (*SSGs*), i.e., *α*-helix, *β*-sheet, or coil-turn (Table 1). We represent a protein of length *l*, as a one-hot encoding *𝓁×* |*S*|-matrix *M*_str_, where *S* = *R × SSG* and *R* is the set of the standard 20 amino acids plus a wildcard *X* for all other amino acids. We order the 63 pairs in *S* lexicographically, first based on the one-letter-codes of the amino acids and then, for a given amino acid, in the order of *α*-helix, *β*-sheet, and coil-turn of the *SSG*. An entry *M*_str_[*i, j*] is 1 if and only if the amino acid at the *i*^*th*^ position belongs to the combination of amino acid and *SSG* at the *j*^*th*^ position in *S*.

**Table 1.** Amino acid-specific secondary structural categories obtained from DSSP and the corresponding secondary structural group (*SSG*) defined in our study.

| Secondary structural category obtained from DSSP | SSG |
| --- | --- |
| α-helix<br>3-helix<br>5-helix | α-helix |
| Residue in isolated β-bridge<br>Extended strand participating in $\beta$ ladder | β-sheet |
| Hydrogen-bonded turn<br>Bend<br>Blank (undetermined output) | coil-turn |

### 2.7. Our deep learning model architecture

Our DL architecture is similar to that of DeepTFactor with some adaptations due to the additional features related to the secondary structural information at each position of the protein (Section 2.6). We keep our DL architecture as similar to DeepTFactor as possible to ensure that the performance difference between StrucTFactor and DeepTFactor could clearly be attributed to the use of secondary structural information, rather than the use of the DL architecture.

Our DL architecture consists of CNNs which process the input matrices of size 1 000 *×* 63. The CNN has three subnetworks (termed *a, b*, and *c*), each consisting of three convolutional layers with an attached batch normalization, a rectified linear unit (ReLU), and a dropout layer with a dropout rate of 0.1. The kernel sizes of the three convolutional layers within the three subnetworks are [4,63], [4,1], and [16,1] for subnetwork *a*, [12,63], [8,1], and [4,1] for subnetwork *b* and [16,63], [4,1], and [4,1] for subnetwork *c*. Each of these layers uses a stride of 1 and a padding of 0. The output of each of the three subnetworks is concatenated into a single feature map, which is processed by a convolutional layer followed by batch normalization, ReLU, max-pooling, and two fully connected layers with batch normalization. The first fully connected layer uses ReLU as an activation function, while the second fully connected layer outputs a single value obtained by a Sigmoid function. Detailed schematics of our deep learning architecture is shown in Supplementary Figure S2. We initialize the learnable parameters in the convolutional layers and fully connected layer by Xavier uniform distribution (Glorot and Bengio, 2010).

### 2.8. Model training and evaluation

We divide the proteins of a dataset into five non-overlapping subsets with stratification. By stratification, we mean that the ratio of the numbers of non-TFs vs. TFs in each subset is the same as the ratio of the numbers of non-TFs vs. TFs in the complete dataset. Given a subset (i.e., 20% of the complete dataset), we use it as an independent test data and divide all of the proteins in the remaining four subsets into training (72% of the complete dataset) and validation (8% of the complete dataset) data. We do this for each of the five subsets and hence obtain five combinations of training, validation, and independent test data.

For the existing methods, i.e., DeepTFactor and DeepReg, for each of the five combinations of training, validation, and test data, we use a constant set of hyperparameter values recommended by the corresponding studies. Similarly, for our proposed method, i.e., StrucTFactor, for each of the five combinations of training, validation, and test data, we use a constant set of hyperparameter values. Because StrucTFactor has a similar deep learning architecture as DeepTFactor, to be as conservative as possible, we use those hyperparameter values for StrucTFactor that were optimized for DeepTFactor. That is, we use a learning rate of 0.001, a batch size of 128, and train a given model for 50 epochs. Considering the three different TF prediction methods (i.e., StrucTFactor, DeepTFactor, and DeepReg), 12 datasets, and five combinations of training, validation, and independent test data, we train and test 3 *×* 12 *×* 5 = 180 TF prediction models.

We evaluate each trained model with respect to the corresponding independent test data using three performance measures: AU-PRC, AU-ROC, and MCC. For MCC, we define a threshold of 0.5, meaning that we predict a protein to be a TF if its TF prediction score is *>* 0.5 and we predict a protein to be non-TF if its TF prediction score is *≤* 0.5. MCC is effective in evaluating model performance on imbalanced datasets because it gives a high score if and only if the prediction is good in all categories (i.e., true positives, false positives, true negatives, and false negatives) of the confusion matrix (Chicco and Jurman, 2020).

### 2.9. Interpretability of trained models

To gain insights into the features influencing the predictions of a trained model, we use our most reliable dataset *D*(*rl, nr*, 3) and apply the integrated gradient method (Sundararajan et al., 2017) from the captum (Kokhlikyan et al., 2020) library of PyTorch (Paszke et al., 2019).

Given a trained model, for a protein of length *𝓁*the input matrix of size *𝓁× q*, where *q* = 21 for DeepTFactor/DeepReg and *q* = 63 for StrucTFactor, the integrated gradient method outputs a matrix of the same size as the input matrix. Each cell of the output matrix indicates the impact of the input feature corresponding to that cell on the predicted result. A score can either be positive or negative, where a positive score means that the input feature at the corresponding cell supports the model to predict TF, while a negative score means that the input feature at the corresponding cell supports the model to predict non-TF. Given the output matrix, we take the sum of each row to get one score per sequence position of the protein.

## 3. Results and discussion

To evaluate whether the use of 3D secondary structural information can more accurately distinguish TFs from non-TFs than just using sequences, we compare our 3D secondary structure-based TF prediction method StrucTFactor with the existing recent state-of-the-art sequence-based TF prediction methods (i.e., DeepTFactor and DeepReg).

First, we compare the performances of the considered methods on the most reliable dataset *D*(*rl, nr*, 3) that has sequence non-redundant proteins and the smallest ratio of the numbers of non-TFs vs. TFs (Section 3.1). Second, to evaluate the effect of different ratios of the numbers of non-TFs vs. TFs (i.e., either 3, 5, or 10) while keeping sequence non-redundancy among proteins, we compare the performances of the methods across three datasets *D*(*rl, nr, s*), where *s* ∈ {3, 5, 10} (Section 3.2). Third, to evaluate the effect of sequence redundancy, we compare the performances of the methods across three sequence redundant datasets *D*(*rl, r, s*) and three sequence non-redundant datasets *D*(*rl, nr, s*) (Section 3.3). Fourth, to evaluate the effect of reliable vs. all predicted 3D protein structures, we compare the performance of StrucTFactor across six datasets *D*(*rl, z, s*) containing reliably predicted 3D protein structures and six datasets *D*(*a, z, s*) containing all predicted 3D protein structures, where *z* ∈ {*nr, r*} (Section 3.4). Fifth, to interpret the prediction performance of StrucTFactor, we look at the parts of proteins that StrucTFactor exploits to correctly distinguish TFs from non-TFs (Section 3.5).

### 3.1. StrucTFactor outperforms existing TF prediction methods on the most confident dataset

For the most reliable dataset *D*(*rl, nr*, 3), StrucTFactor clearly outperforms DeepTFactor and DeepReg with respect to all three performance measures (Figure 3).

**Figure 3:**
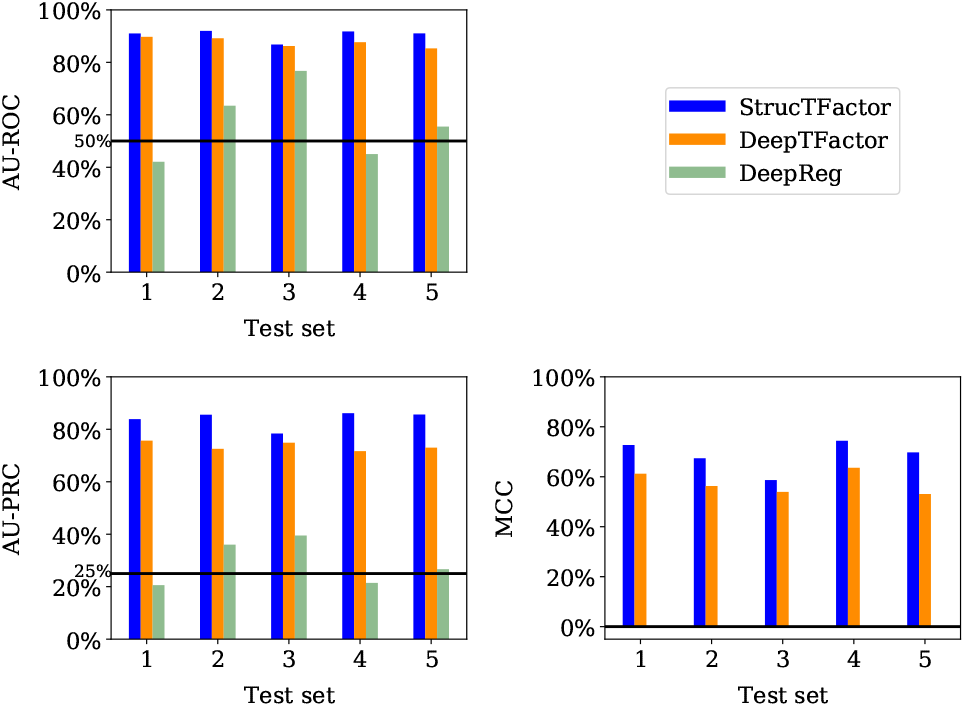
Performances of StrucTFactor (blue), DeepTFactor (orange), and DeepReg (green) based on AU-ROC (upper left panel), AU-PRC (lower left panel) and MCC (lower right panel) for each of the five test sets corresponding to *D*(*rl, nr*, 3). The horizontal lines in the upper and lower left panels indicate the performance values for AU-ROC (50%) and AU-PRC (25%), respectively, that are expected by random chance. DeepReg predicts all proteins as non-TFs, which results in undefined MCC scores.

Interestingly, while both StrucTFactor and DeepTFactor perform significantly better than random with AU-ROC values well above 50%, DeepReg’s performance is close to random for most test sets (Figure 3, upper left panel). This result could be due to the small size of *D*(*rl, nr*, 3), which likely does not suffice to train the large number of parameters of the DeepReg architecture. With respect to AU-PRC, StrucTFactor achieves scores from *∼*78% to *∼*86%, outperforming DeepTFactor by a margin between *∼*4% and *∼*14%, depending on the test set (Figure 3 and Supplementary Table S3). With respect to MCC, StrucTFactor achieves scores from *∼*58% to *∼*74%, outperforming DeepTFactor by a margin between *∼*5% and *∼*17%, depending on the test set (Figure 3). These results signify that using 3D secondary structural information (as in StrucTFactor), instead of just the sequence information (as in DeepTFactor or DeepReg), can more accurately distinguish TFs from non-TFs.

Note that the performances of both DeepTFactor and DeepReg were shown to be much higher (e.g., *∼*94% MCC for DeepTFactor) in their original studies than what we find in our evaluation in this subsection. This is because both DeepTFactor and DeepReg were evaluated on sequence redundant datasets, leading to possible data leakage across training and test sets which likely resulted in their overestimated performance scores (Bernett et al., 2024). In contrast, our evaluations in this subsection are based on a sequence non-redundant dataset that prevents data leakage and thus provide a more accurate estimation of a method’s performance.

### 3.2. StrucTFactor outperforms existing TF prediction methods irrespective of the class assignment ratio

We evaluate the performances of the considered methods across different class assignment ratios based on three datasets *D*(*rl, nr, s*), with *s* ∈ {3, 5, 10}. While both StrucTFactor and DeepTFactor perform significantly better than random with AU-ROC values well above 50%, DeepReg’s performance is close to random for two out of the three datasets (Figure 4).

**Figure 4:**
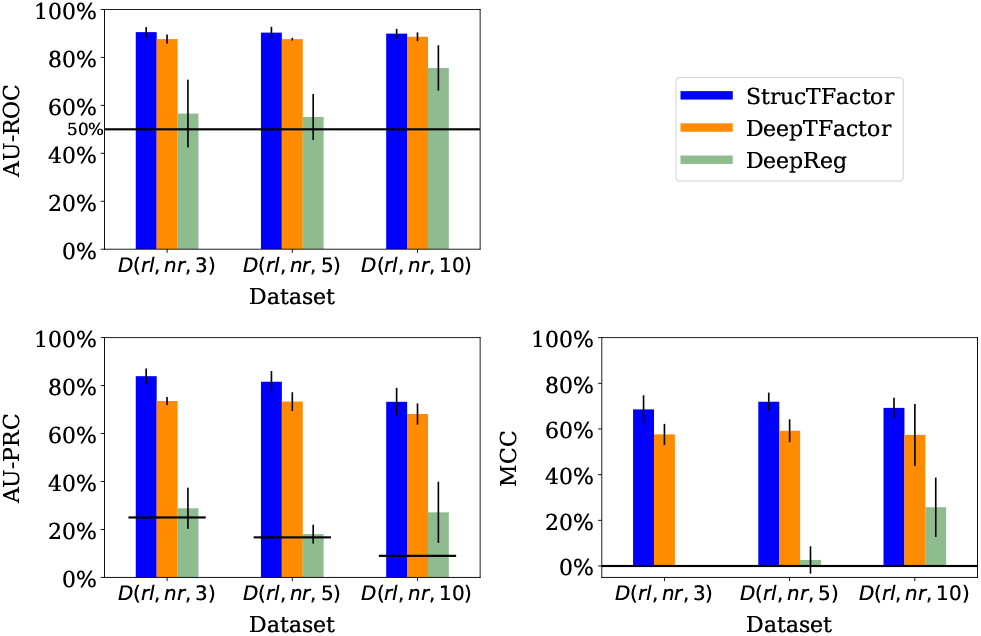
Performances of StrucTFactor (blue), DeepTFactor (orange), and DeepReg (green) based on AU-ROC (upper left panel), AU-PRC (lower left panel) and MCC (lower right panel) for *D*(*rl, nr, s*), where *s* ∈ {3, 5, 10}. The height of a bar represents the average performance over the five test sets, while the vertical line on a bar represents the corresponding standard deviation. The horizontal lines in the upper and lower left panels indicate the performance values for AU-ROC (50%) and AU-PRC, respectively, that are expected by random chance. Because the AU-PRC random chance performance value for a dataset is the fraction of TFs in the dataset, it is individually defined for each dataset.

With respect to AU-PRC, the performances of both StrucTFactor and DeepTFactor slightly decrease with an increase in the ratio of the numbers of non-TFs vs. TFs (Figure 4). Both methods perform the best for *D*(*rl, nr*, 3), the second best for *D*(*rl, nr*, 5), and the worst for *D*(*rl, nr*, 10). However, with respect to MCC, both methods perform slightly better for *D*(*rl, nr*, 5) compared to *D*(*rl, nr*, 3) and *D*(*rl, nr*, 10). For example, the MCC values (averaged over the five test sets) for StrucTFactor are *∼*68.6%, *∼*71.9%, and *∼*69.2% for *D*(*rl, nr*, 3), *D*(*rl, nr*, 5), and *D*(*rl, nr*, 10), respectively. The above results could be due to the trade-off between the class assignment ratio and the number of proteins in a dataset, both of which can affect the training of an ML model. While *D*(*rl, nr*, 3) has the smallest ratio between the numbers of non-TFs vs. TFs, it also has the least number (2 540) of proteins on which an ML model can be trained. The larger ratio of the numbers of non-TFs vs. TFs in *D*(*rl, nr*, 5) compared to *D*(*rl, nr*, 3) leads to a considerably larger number (3 810) of proteins to train an ML model. However, for *D*(*rl, nr*, 10), although there is a higher number (6 985) of proteins, the performance decreases. This may be due to the fact that the ratio of the numbers of non-TFs vs. TFs is a bit too large, which can negatively affect the training of the ML model such that it is not able to generalize.

Unlike StrucTFactor and DeepTFactor, DeepReg’s performance for AU-ROC and MCC increases with an increase in the ratio of the numbers of non-TFs vs. TFs. This could mean that DeepReg’s performance might increase for a larger ratio of the numbers of non-TFs vs. TFs. To verify this hypothesis, we ran all three methods on all data from the reliable sequence non-redundant set of proteins containing 20 249 non-TFs and 635 TFs (non-TF vs. TF ratio of 32). We find that our hypothesis does not hold (Supplementary Table S4). The performance of DeepReg for this dataset is worse than its performance for *D*(*rl, nr*, 10) and it is inferior to the other two methods.

Irrespective of the class assignment ratio across the three datasets, StrucTFactor consistently outperforms DeepTFactor (Figure 4). Specifically, with respect to AU-PRC (averaged over the five test sets), StrucTFactor improves upon DeepTFactor by a margin between *∼*5% and *∼*10%, depending on the dataset. With respect to MCC (averaged over the five test sets), StrucTFactor improves upon DeepTFactor by a margin between *∼*11% and *∼*13%, depending on the dataset. To quantify whether the performance of StrucTFactor is significantly better than DeepTFactor, given a performance measure, we compare their performance values over all 15 combinations (three datasets with five test sets each) using the paired Wilcoxon signed-rank test (Woolson, 2008). We find that StrucTFactor significantly (*p*-value *<* 0.005) outperforms DeepTFactor with respect to each of the three performance measures.

### 3.3. StrucTFactor outperforms existing TF prediction methods irrespective of sequence redundancy

To study the effect of sequence redundancy on the performance of a method, we compare the results for three sequence non-redundant datasets *D*(*rl, nr, s*) with the results for three sequence redundant datasets *D*(*rl, r, s*), with *s* ∈ {3, 5, 10}. Because high sequence identities across proteins in the training and test data can artificially boost the performances of trained ML models (Hou et al., 2022), we expect to see an increase in the performance of a method on sequence redundant datasets in comparison to the corresponding sequence non-redundant datasets. This is exactly what we find (Table 2). For example, DeepTFactor achieves an average MCC of *∼*58% over the five test sets of *D*(*rl, nr*, 3) (a sequence non-redundant dataset), while it achieves an average MCC of *∼*97% over the five test sets of the corresponding sequence redundant dataset *D*(*rl, r*, 3). For sequence redundant datasets, DeepTFactor and DeepReg already achieve performances close to 100%. We note that in both DeepTFactor and DeepReg studies (Kim et al., 2021; Ledesma-Dominguez et al., 2024) *only* the sequence redundant datasets were used to train and test their respective models, which resulted in their boosted performances that we also verify in our study. Nevertheless, even in the small scope for performance improvements for sequence redundant datasets, StrucTFactor consistently outperforms DeepTFactor and DeepReg based on all three performance measures (Table 2 and Supplementary Table S5).

**Table 2.** Influence of sequence redundancy on TF prediction performance of the considered methods. For *D*(*rl, z*, 3), we show average (over the five test sets) AU-PRC, MCC, and AU-ROC values for the sequence non-redundant (*z* = *nr*) and sequence redundant (*z* = *r*) datasets. As DeepReg predicts all proteins as non-TFs for *D*(*rl, nr*, 3), the corresponding MCC score is undefined which we denote by *nan*. The best result for each performance measure and sequence redundancy choice is in bold font. The results are for datasets with the numbers of non-TFs vs. TFs ratio of 3. The corresponding results for the alternative ratios of 5 and 10 are qualitatively similar (Supplementary Table S5).

|  | AU-PRC |  | MCC |  | AU-ROC |  |
| --- | --- | --- | --- | --- | --- | --- |
| | $z = nr$ | $z = r$ | $z = nr$ | $z = r$ | $z = nr$ | $z = r$ |
| StrucTFactor | **0.839** | **0.988** | **0.686** | **0.975** | **0.905** | **0.993** |
| DeepTFactor | 0.736 | 0.986 | 0.576 | 0.965 | 0.876 | 0.991 |
| DeepReg | 0.288 | 0.976 | nan | 0.878 | 0.566 | 0.988 |

### 3.4. StrucTFactor is robust across different 3D protein structural quality

To investigate the influence of reliability of the predicted 3D protein structures on the performance of StrucTFactor, we compare its results for the six datasets from reliableAF proteins*D*(*rl, z, s*) with its results for the six datasets from allAF proteins *D*(*a, z, s*), where *z* ∈ {*nr, r*} and *s* ∈ {3, 5, 10}.

Although allAF datasets contain many 3D protein structures with confidence scores lower than the confidence score that we use to identify reliable 3D protein structures (Section 2.2), StrucTFactor performs better (or comparable) for allAF datasets than for the corresponding reliableAF datasets (Table 3). Additionally, similar to the reliableAF datasets (see previous subsections of results), StrucTFactor outperforms DeepTFactor and DeepReg on allAF datasets (Table 3), although with a smaller margin compared to the reliableAF datasets.

**Table 3.** Influence of the quality of the predicted 3D protein structures on the performance of StrucTFactor. For *D*(*x, nr*, 3), we show average (over the five test sets) AU-PRC, MCC, and AU-ROC values for reliableAF (*x* = *rl*) and allAF (*x* = *a*). The best result for each performance measure and protein 3D structure reliability choice is in bold font. The results are for the datasets with the numbers of non-TFs vs. TFs ratio of 3. The corresponding results for ratios of 5 and 10 are qualitatively similar (Supplementary Table S5).

|  | AU-PRC |  | MCC |  | AU-ROC |  |
| --- | --- | --- | --- | --- | --- | --- |
| | $x = rl$ | $x = a$ | $x = rl$ | $x = a$ | $x = rl$ | $x = a$ |
| StrucTFactor | **0.839** | **0.860** | **0.686** | **0.739** | **0.905** | **0.911** |
| DeepTFactor | 0.736 | 0.835 | 0.576 | 0.685 | 0.876 | 0.904 |
| DeepReg | 0.288 | 0.776 | nan | 0.542 | 0.566 | 0.887 |

The above results could be due to the following reasons. The number of proteins in allAF datasets is larger than the number of proteins in the corresponding reliableAF datasets (Figure 2). This could improve the training of an ML model. Thus, the performances of each method are, in general, higher for allAF datasets than for the corresponding reliableAF datasets. However, the additional proteins in the allAF datasets are of lower 3D structural quality. Because StrucTFactor (unlike DeepTFactor and DeepReg) relies on 3D protein structures, the increase in the number of proteins in the allAF datasets compared to the corresponding reliableAF datasets does not boost the performance of StrucTFactor as much as it does for DeepTFactor and DeepReg. Nevertheless, to statistically quantify the superiority of StrucTFactor over DeepTFactor or DeepReg, we compare their performance values over all 30 combinations (six allAF datasets with five test sets each), using the paired Wilcoxon signed-rank test. We find that StrucTFactor statistically significantly (*p*-value *<* 0.03) outperforms both DeepTFactor and DeepReg with respect to each of the three performance measures.

### 3.5. StrucTFactor captures DNA-binding domain regions of transcription factors

To get insights into why StrucTFactor performs well on the TF prediction task, we characterize the specific regions of a protein that StrucTFactor focuses on when making predictions. We then compare these regions only to those identified by DeepTFactor, given that DeepTFactor performs much better than DeepReg in our evaluations. Given a protein, we use the concept of integrated gradients that gives a score to each sequence position (Section 2.9).

We do this for each of the 395 TFs from the most reliable dataset, i.e., *D*(*rl, nr*, 3), for which there is DBD information (i.e., sequence positions of a protein that form a DBD) in UniProt. Given the DBD information of a TF, we take the sum of the integrated gradient scores of all sequence positions in the DBD region as the TF’s DBD-related score and the sum of the integrated gradient scores of all sequence positions not present in the DBD region as the TF’s non-DBD-related score.

We find that the DBD-related scores of TFs predicted by StrucTFactor are mostly positive and are in general higher than the DBD-related scores of TFs predicted by DeepTFactor (Figure 5). This signifies that StrucTFactor has, in comparison to DeepTFactor, a larger focus on the known DBD regions of TFs. Additionally, the difference between the DBD-related scores and non-DBD-related scores are significantly larger for those predicted by StrucTFactor than for those predicted by DeepTFactor (Figure 5 and Supplementary Figure S3). This signifies that the existence of DBDs in TFs has a larger influence on their positive predictions by StrucTFactor than by DeepTFactor. Note that we do not use the DBD information while training any model. Nevertheless, it seems that the models can inherently capture such DBD information during the process of training.

**Figure 5:**
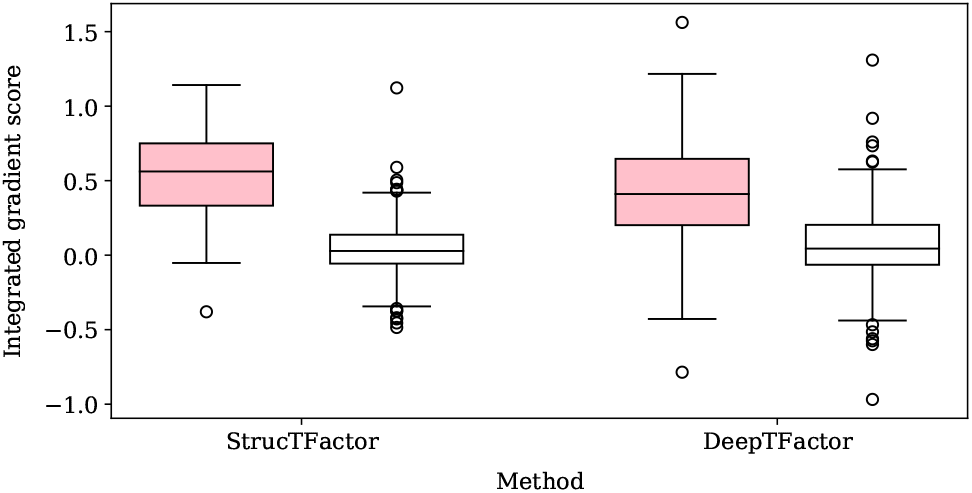
Distribution of DBD-related (pink) and non-DBD-related (white) integrated gradient scores of all 395 TFs from the *D*(*rl, nr*, 3) dataset that has DBD information in UniProt.

## 4. Conclusion

We propose a novel data-driven DL-based TF prediction method called StrucTFactor, which is the first of its kind to use 3D secondary structural features of proteins. Our comprehensive evaluation shows that StrucTFactor significantly outperforms the existing state-of-the-art TF prediction methods that rely only on protein sequences.

We develop DL models using all reviewed protein sequences from UniProt. These originate from many different species. As in the evaluation of previous TF prediction methods, we define the ground truth TFs based on GO annotations. We note that using GO annotations to define TFs might not be ideal. For example, such annotations could be biased towards homologous proteins (Gaudet and Dessimoz, 2017), which could limit the variety of TFs that a model can be trained on, and thus limits the trained model’s generalizability. To avoid this, we create datasets with no potential homologs by including only proteins such that any pairwise identity is less than 30%. We also note that a species level analysis, rather than an analysis over all species combined (our approach), could provide additional biological insights. For example, it has been shown that 3D structural characteristics of TFs vary across species of different complexity (Yruela et al., 2017). Additionally, a species level analysis inherently avoids inclusion of orthologs, but not necessarily paralogs. To potentially remove paralogs, an approach similar to what we already use in this work should be used. As is, a species-level analysis entails a significant effort and is out of the scope of this work.

To fully understand the transcriptional regulatory role of a protein, it is not sufficient to just predict whether a protein is a TF or not. Additionally, the understanding would require further identifying the DBDs, the genomic binding sites, and the regulated genes corresponding to a TF, which is typically done by experimental methods, such as DNA binding and knockdown or over-expression of the protein in question (Lambert et al., 2018). Such experimental methods are costly. Because biomolecular processes, e.g., alternative splicing and post-translational modifications, create a variety of protein forms (Smith et al., 2021), experimentally testing each protein form regarding its ability to act as a TF is not feasible. In this regard, computational methods for TF predictions (e.g., StrucTFactor) could be used as a first step to reduce the search space of proteins for further experimental validations.

StrucTFactor is especially applicable to identify those novel TFs that do not show sequence similarity to any of the DBDs of known TFs. Interestingly, despite not using any DBD information in the training process, StrucTFactor seems to capture known DBD information better than the considered existing TF prediction methods. As is, StrucTFactor can not only predict novel TFs, but could also be used to identify DBDs of a novel TF and could thus help in better characterizing the transcriptional regulatory role of the TF. However, further analyses are required to verify the latter, which is out of the scope of this work.

Overall, our study highlights the importance of considering 3D secondary structural features of proteins for solving the TF prediction task. We emphasize that we train our models based on predicted 3D structures of proteins. This shows that the recent advancements in 3D protein structural predictions can be utilized to considerably improve the annotation of TFs. Future research focusing on protein features based on 3D tertiary structures of proteins has the potential to further improve the prediction of TFs.

## 6. Author contributions statement

Conceived the study: F.N. and K.N. Collected and processed the data: F.N. and J.L. Designed the methodology: F.N. and K.N. Designed the experiments: F.N., J.L., and K.N. Performed the experiments: F.N. and J.L. Analyzed the results: F.N., J.L., J.G., J.B., and K.N. Wrote the paper: F.N., J.L., S.K., and K.N. Read and approved the paper: F.N., J.L., J.G., S.K., J.B., and K.N. Supervised the study: K.N.

