## Supplementary materials for "Transcription factor prediction using protein 3D secondary structures"

### Supplementary materials: Transcription factor prediction using protein 3D secondary structures

[1]Institute for Computational Systems Biology, Universität Hamburg, Albert-Einstein-Ring 8-10, 22761 Hamburg, Germany, [2]Faculty of Mathematics, Informatics and Natural Sciences, ZBH - Center for Bioinformatics, Universität Hamburg, Albert-Einstein-Ring 8-10, 22761 Hamburg, Germany, [3]Department of Mathematics and Computer Science, University of Southern Denmark, Campusvej 55, 5230 Odense, Denmark and [4]Center for Data and Computing in Natural Sciences, Universität Hamburg, Albert-Einstein-Ring 8-10, 22761 Hamburg, Germany

[*]Corresponding author, [†]Equal contribution

#### I Supplementary sections

##### S1. Information regarding 3D secondary structural data from PDB

To evaluate the possibility of using experimentally determined structures from the Protein Data Bank (PDB) (Berman et al., 2000), we analyze the PDB data as follows. We consider all UniProt proteins with PDB entries having 3D resolutions of $\leq 3\mathring{A}$, resulting in 25 442 proteins. Because one protein can have multiple PDB chain entries, for each of the 25 442 proteins, we select the PDB chain entry with the highest number of 3D resolved residues (i.e., the PDB chain with the maximum number of residues with secondary structural information). Then, for each protein, to quantify the fraction of residues with secondary structural information (named "3D residue coverage"), we divide the number of 3D structurally resolved residues of a protein with its total number of residues.

We find that most proteins have $\leq 0.5$ 3D residue coverage (Supplementary Figure S1). Because the focus of this study is to build a TF prediction model, we need sufficient number of TFs to train such a model. We find that, among the 25 442 proteins, there are 1 147 TFs. However, only 295 out of the 1 147 (i.e., ~26%) TFs have a 3D residue coverage of $\geq 0.5$, and this percentage decreases drastically with an increase in the 3D residue coverage value (Supplementary Table S1). This analysis shows the limited availability of the PDB data to build a 3D structure-based machine learning model for TF prediction.

##### S2. Datasets based on allAF proteins

Similar to reliableAF (Section 2.5 in the main paper), we create six datasets, i.e., $D(a, z, s)$ where $z \in \{nr, r\}$ and $s \in \{3, 5, 10\}$, for allAF. Out of all 524 674 allAF proteins, there are 19 014 TFs and 505 660 non-TFs. To remove the class imbalance problem, we take three random samples from all 505 660 non-TFs, such that the number of non-TFs is either 3, 5, or 10 times more than the number of TFs, resulting in three datasets $D(a, r, 3)$, $D(a, r, 5)$, and $D(a, r, 10)$. Additionally, we remove sequence redundancy among the 524 674 proteins in the similar manner as we do for reliableAF, which results in 61 025 (with 3 798 TFs and 57 227 non-TFs) proteins. Then, we remove the class imbalance problem as above from the dataset with sequence non-redundant proteins, resulting in three datasets named $D(a, nr, 3)$, $D(a, nr, 5)$, and $D(a, nr, 10)$, see Figure 2 in the main paper.

#### II Supplementary figures

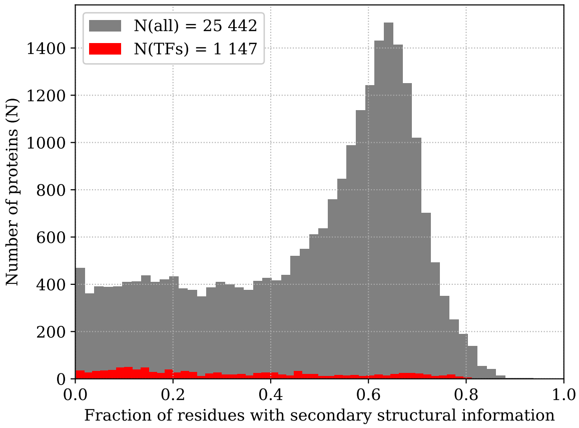


Figure S1: Distribution of fractions of protein residues with secondary structural information (i.e., 3D residue coverage) from PDB. The grey bars represent the distribution of 3D residue coverage for all 25 442 UniProt proteins with a PDB entry having the 3D resolution of $\leq 3\AA$. The red bars show the subset of 25 442 (i.e., 1 147) proteins that are TFs.

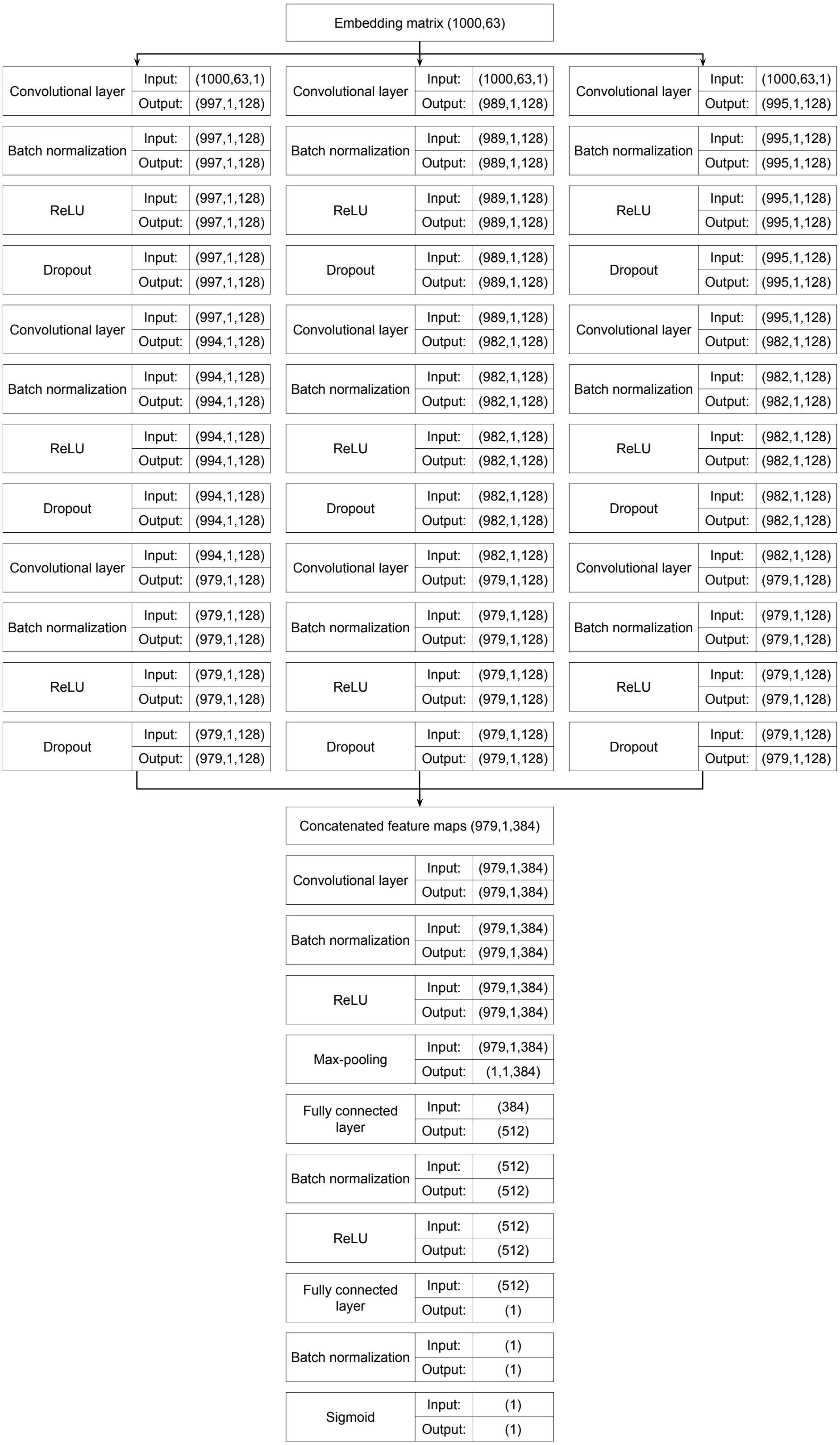


Figure S2: Deep learning architecture of StrucTFactor. Figure adapted from (Kim et al., 2021).

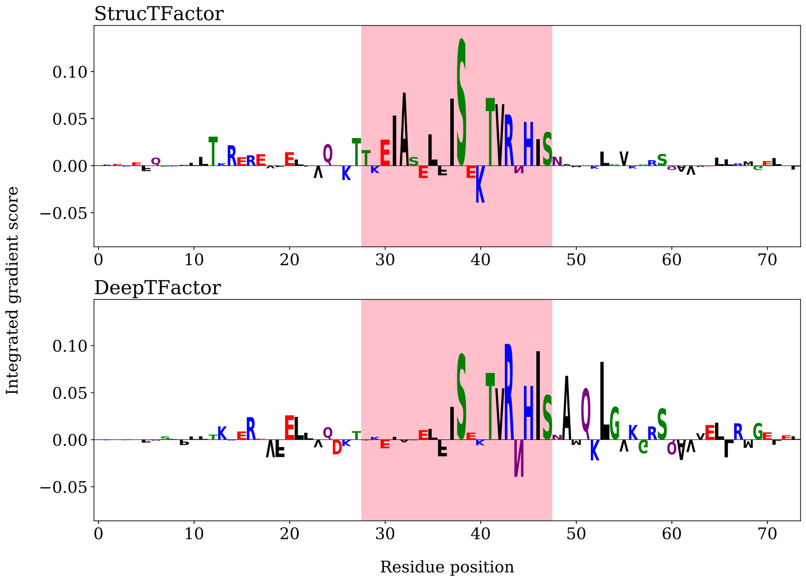


Figure S3: Integrated gradient score of each sequence position of the Spore germination protein GerE (UniProt ID *P11470*). This protein achieves the highest TF prediction score by both StrucTFactor and DeepTFactor. We highlight the known DBD region in pink (ground truth knowledge and not part of the predictions). For each position along the protein sequence on the $X$-axes, the corresponding integrated gradient score is represented on the $Y$-axes, quantified by the size and orientation of the amino acid letter.

#### III Supplementary tables

**Table S1** Fraction of residues in a protein with secondary structural information. We show results for all considered PDB proteins and all considered PDB TFs.

| Fraction of residues in a protein with secondary structural information $\geq$ x | x = 0.0 | x = 0.5 | x = 0.6 | x = 0.7 | x = 0.8 | x = 0.9 |
|---|---|---|---|---|---|---|
| Number of all considered UniProt proteins with sufficient 3D structural resolution | 25 442 | 14 439 | 9 747 | 2 664 | 295 | 10 |
| Number of all considered UniProt TFs with sufficient 3D structural resolution | 1 147 | 295 | 217 | 91 | 5 | 0 |

**Table S2** Summary of the GO terms used for labeling proteins as TFs versus non-TFs, adopted from (Kim et al., 2021) and (Ledesma-Dominguez et al., 2024).

| Type | GO Terms | Description |
|---|---|---|
| Transcription factor | GO:0000976 | Transcription regulatory region sequence-specific DNA binding |
| Transcription factor | GO:0000977 | RNA polymerase II transcription regulatory region sequence-specific DNA binding |
| Transcription factor | GO:0000978 | RNA polymerase II cis-regulatory region sequence-specific DNA binding |
| Transcription factor | GO:0000979 | RNA polymerase II core promoter sequence-specific DNA binding |
| Transcription factor | GO:0000981 | DNA-binding Transcription factor activity, RNA polymerase II-specific |
| Transcription factor | GO:0000984 | Bacterial-type RNA polymerase transcription regulatory region sequence-specific DNA binding |
| Transcription factor | GO:0000985 | Bacterial-type RNA polymerase core promoter sequence-specific DNA binding |
| Transcription factor | GO:0000986 | Bacterial-type cis-regulatory region sequence-specific DNA binding |
| Transcription factor | GO:0000987 | cis-regulatory region sequence-specific DNA binding |
| Transcription factor | GO:0000992 | RNA polymerase III cis-regulatory region sequence-specific DNA binding |
| Transcription factor | GO:0000995 | RNA polymerase III general transcription initiation factor activity |
| Transcription factor | GO:0001046 | Core promoter sequence-specific DNA binding |
| Transcription factor | GO:0001163 | RNA polymerase I transcription regulatory region sequence-specific DNA binding |
| Transcription factor | GO:0001164 | RNA polymerase I core promoter sequence-specific DNA binding |
| Transcription factor | GO:0001165 | RNA polymerase I cis-regulatory region sequence-specific DNA binding |
| Transcription factor | GO:0001216 | DNA-binding transcription activator activity |
| Transcription factor | GO:0001227 | DNA-binding transcription repressor activity, RNA polymerase II-specific |
| Transcription factor | GO:0003700 | DNA-binding TF activity |
| Transcription factor | GO:0034246 | Mitochondrial sequence-specific DNA-binding TF activity |
| Transcription factor | GO:0098531 | Ligand-activated TF activity |
| Transcription factor | GO:0106250 | DNA-binding transcription repressor activity, RNA polymerase III-specific |
| Transcription regulation | GO:0001228 | DNA-binding transcription activator activity, RNA polymerase II-specific |
| Transcription regulation | GO:0006351 | Transcription, DNA-templated |
| Transcription regulation | GO:0006355 | Regulation of transcription, DNA-templated |
| Transcription regulation | GO:0043433 | Negative regulation of DNA-binding TF activity |
| Transcription regulation | GO:0045892 | Negative regulation of transcription, DNA-templated |
| Transcription regulation | GO:0045893 | Positive regulation of transcription, DNA-templated |
| Transcription regulation | GO:0051090 | Regulation of DNA-binding TF activity |
| Transcription regulation | GO:0051091 | Positive regulation of DNA-binding TF activity |
| Transcription regulation | GO:2000142 | Regulation of DNA-templated transcription, initiation |
| Transcription regulation | GO:2000143 | Negative regulation of DNA-templated transcription, initiation |
| Transcription regulation | GO:2000144 | Positive regulation of DNA-templated transcription, initiation |
| DNA binding | GO:0003677 | DNA binding |
| DNA binding | GO:0008301 | DNA binding, bending |
| DNA binding | GO:0043565 | Sequence-specific DNA binding |
| DNA binding | GO:0050692 | DNA binding domain binding |

**Table S3** Performances of StrucTFactor, DeepTFactor, and DeepReg on the dataset $D(rl, nr, 3)$. For each method, the results are shown for each of the five test sets individually as well as for the average of the five test sets (last column named *Mean*). The best performing method for a given test set or for the average of the five test sets is in bold font. DeepReg predicts all proteins as non-TFs, which results in undefined ("nan") MCC scores.

| | AU-PRC | | | | | |
|---|---|---|---|---|---|---|
| CV-Fold | 1 | 2 | 3 | 4 | 5 | Mean |
| StrucTFactor | **0.8388** | **0.8556** | **0.7839** | **0.8613** | **0.8564** | **0.8392** |
| DeepTFactor | 0.7567 | 0.7256 | 0.7489 | 0.7166 | 0.7301 | 0.7356 |
| DeepReg | 0.2057 | 0.3602 | 0.3951 | 0.2144 | 0.2666 | 0.2884 |
| | MCC | | | | | |
| CV-Fold | 1 | 2 | 3 | 4 | 5 | Mean |
| StrucTFactor | **0.7270** | **0.6739** | **0.5866** | **0.7440** | **0.6975** | **0.6858** |
| DeepTFactor | 0.6122 | 0.5630 | 0.5395 | 0.6360 | 0.5308 | 0.5763 |
| DeepReg | nan | nan | nan | nan | nan | nan |
| | AU-ROC | | | | | |
| CV-Fold | 1 | 2 | 3 | 4 | 5 | Mean |
| StrucTFactor | **0.9101** | **0.9198** | **0.8678** | **0.9176** | **0.9102** | **0.9051** |
| DeepTFactor | 0.8971 | 0.8917 | 0.8621 | 0.8768 | 0.8530 | 0.8761 |
| DeepReg | 0.4209 | 0.6346 | 0.7673 | 0.4502 | 0.5550 | 0.5656 |

**Table S4** Performances of StrucTFactor, DeepTFactor, and DeepReg on all data from the reliable non-redundant set of proteins containing 20,249 non-TFs and 635 TFs. Given a dataset, the results are shown as the average of the five test sets. The best average performance value corresponding to each metric is in bold font.

| | AU-PRC | MCC | AU-ROC |
|---|---|---|---|
| StrucTFactor | **0.5490** | **0.5654** | **0.8262** |
| DeepTFactor | 0.4641 | 0.5207 | 0.7536 |
| DeepReg | 0.1000 | 0.1658 | 0.6077 |

**Table S5** Variation in the performances of StrucTFactor, DeepTFactor, and DeepReg on different sequence redundancies and class assignment ratios for our 12 datasets $D(x, z, s)$ where $x \in \{rl, a\}$, $z \in \{nr, r\}$, and $s \in \{3, 5, 10\}$ (see Section 2.5 and Figure 2 in the main paper for details). Given a dataset, the results are shown as average of the five test sets. Given a dataset and a performance measure, the best method performance is in bold font.

| $D(x,z,s)$ | $x=rl$ $z=nr$ $s=3$ | $x=rl$ $z=nr$ $s=5$ | $x=rl$ $z=nr$ $s=10$ | $x=rl$ $z=r$ $s=3$ | $x=rl$ $z=r$ $s=5$ | $x=rl$ $z=r$ $s=10$ | $x=a$ $z=nr$ $s=3$ | $x=a$ $z=nr$ $s=5$ | $x=a$ $z=nr$ $s=10$ | $x=a$ $z=r$ $s=3$ | $x=a$ $z=r$ $s=5$ | $x=a$ $z=r$ $s=10$ |
|---|---|---|---|---|---|---|---|---|---|---|---|---|
| | AU-PRC | | | | | | | | | | | |
| StrucTFactor | **0.8392** | **0.8158** | **0.7322** | **0.9876** | **0.9890** | **0.9844** | **0.8601** | **0.8225** | **0.7892** | **0.9815** | **0.9766** | 0.9691 |
| DeepTFactor | 0.7356 | 0.7332 | 0.6815 | 0.9858 | 0.9869 | 0.9811 | 0.8354 | 0.8118 | 0.7449 | 0.9784 | 0.9745 | **0.9702** |
| DeepReg | 0.2884 | 0.1806 | 0.2715 | 0.9755 | 0.9426 | 0.9708 | 0.7764 | 0.7125 | 0.6573 | 0.9684 | 0.9622 | 0.9505 |
| | MCC | | | | | | | | | | | |
| StrucTFactor | **0.6858** | **0.7190** | **0.6923** | **0.9747** | **0.9737** | **0.9718** | **0.7391** | **0.7394** | **0.7527** | **0.9514** | **0.9477** | **0.9484** |
| DeepTFactor | 0.5763 | 0.5922 | 0.5741 | 0.9653 | 0.9694 | 0.9658 | 0.6850 | 0.7102 | 0.7075 | 0.9381 | 0.9439 | 0.9434 |
| DeepReg | nan | 0.0264 | 0.2574 | 0.8775 | 0.8507 | 0.9108 | 0.5417 | 0.5561 | 0.5950 | 0.8905 | 0.8826 | 0.9066 |
| | AU-ROC | | | | | | | | | | | |
| StrucTFactor | **0.9051** | **0.9030** | **0.8988** | **0.9925** | **0.9941** | **0.9920** | **0.9114** | **0.9121** | **0.9069** | **0.9883** | **0.9880** | 0.9829 |
| DeepTFactor | 0.8761 | 0.8761 | 0.8859 | 0.9914 | 0.9921 | 0.9908 | 0.9043 | 0.9028 | 0.8852 | 0.9870 | 0.9866 | **0.9865** |
| DeepReg | 0.5656 | 0.5516 | 0.7559 | 0.9877 | 0.9820 | 0.9900 | 0.8873 | 0.8838 | 0.8969 | 0.9835 | 0.9856 | 0.9850 |
